# Drug Response Pharmacogenetics for 200,000 UK Biobank Participants

**DOI:** 10.1101/2020.08.09.243311

**Authors:** Gregory McInnes, Russ B Altman

## Abstract

Pharmacogenetics studies how genetic variation leads to variability in drug response. Guidelines for selecting the right drug and right dose to patients based on their genetics are clinically effective, but are still widely unused. For some drugs, the normal clinical decision making process may lead to the optimal dose of a drug that minimizes side effects and maximizes effectiveness. Without measurements of genotype, physicians and patients may observe and adjust dosage in a manner that reflects the underlying genetics. The emergence of genetic data linked to longitudinal clinical data in large biobanks offers an opportunity to confirm known pharmacogenetic interactions as well as discover novel associations by investigating outcomes from normal clinical practice. Here we use the UK Biobank to search for pharmacogenetic interactions among 200 drugs and 9 genes among 200,000 participants. We identify associations between pharmacogene phenotypes and drug maintenance dose as well as side effect incidence. We find support for several known drug-gene associations as well as novel pharmacogenetic interactions.

## Introduction

Pharmacogenetics promises to revolutionize patient care by offering personalized drug selection and dosage based on an individual’s genetics^1^. Variations in genes that encode proteins involved in drug pharmacokinetics and pharmacodynamics lead to interindividual heterogeneity in drug response and can greatly affect clinical outcome. Dosage guidelines have been developed by organizations such as the Clinical Implementation of Pharmacogenetics Consortium (CPIC; cpicpgx.org) to aid physicians in incorporating pharmacogenetics into their practice, however the adoption of pharmacogenetics by practicing physicians has been slow^2,3^.

Doctor’s may not directly be using pharmacogenetics to inform practice, but genetics influences how patients respond to drugs nonetheless. Some drugs, such as warfarin, have a narrow therapeutic index and blood concentration of the drug must be frequently measured to ensure patient safety^4^. The ultimate dose at which the patient achieves the appropriate, stable, blood concentration of the drug is the maintenance dose. For warfarin, this dose is strongly influenced by genetic factors such including variations in the metabolizing enzymes CYP2C9 and CYP4F2, as well as the drug target VKORC1.

In other instances genetic variation may lead patients to be at higher risk for side effects. The frequently prescribed drug simvastatin has well known pharmacogenetic interactions with *SLCO1B1* that can lead to simvastatin-induced myopathy^5^. Individuals with poor functioning SLCO1B1 are at higher risk for simvastatin-induced myopathy. CPIC guidelines for simvastatin recommend that individuals with poor functioning SLCO1B1 take a reduced simvastatin dose or a different drug altogether.

Numerous pharmacogenetic drug-gene relationships have been discovered, but most pharmacogenetic studies are small and narrowly focused. The use of electronic health record and biobank scale data as a means for pharmacogenetic discovery and validation of known relationships has been proposed, but until recently databases linking clinical data with genetic data for a large number of patients were unavailable^1,6^. Biobanks offer an opportunity to retrospectively assess known drug-gene relationships in a clinical setting as well as offer the opportunity to discover new drug-gene associations. Biobanks and electronic health records have been used to perform targeted association studies between genomics and response to individual drugs^7^ as well as characterize frequency of pharmacogenetic alleles in populations^8,9^, but studies of drug response across a large number of drugs have not yet been performed.

The UK Biobank has now been widely used to perform genome-wide association studies on a wide variety of traits, but it also includes primary care data from the United Kingdom’s National Health System^10^. This dataset offers longitudinal, structured clinical data for more than 220,000 participants that includes diagnoses, laboratory tests, and prescription data. This dataset offers a unique opportunity to identify associations between drug response phenotypes and genetics. Here we present a retrospective pharmacogenetic analysis linking drug exposure for 200 drugs to clinical outcome using the UK Biobank primary care data. We focus on two types of clinical outcomes of interest: maintenance dose and side effect incidence.

## Methods

### Pharmacogenetic Allele Calling

We investigated drug-gene relationships for nine important pharmacogenes in the UK Biobank for 222,114 participants using primary care data from the National Health System, provided by the UK Biobank^10^. The pharmacogenetic alleles used in this study were derived from a previously reported procedure, described here in brief^8^. The genetic data used in this study was imputed from the Axiom Biobank Array for each participant. We included nine genes in our analysis: *CYP2B6, CYP2C19, CYP2C9, CYP2D6, CYP3A5, CYP4F2, SLCO1B1, TPMT*, and *UGT1A1*. The proteins encoded by these genes play critical roles in drug pharmacokinetics and each is included in a CPIC dosing guideline for a drug. We assigned pharmacogenetic phenotypes for each gene using PGxPOP (https://github.com/PharmGKB/PGxPOP). The analysis was limited to individuals of European descent, defined as participants who self-reported as European and were confirmed as European using principal component analysis.

### Drug Dosage Association with Pharmacogenetics

Drugs used in this study were derived from the PharmGKB curated drug list (https://www.pharmgkb.org/downloads, drugs.zip)^11^. For each drug, we extracted prescription information from the UK Biobank primary care prescription data by matching the drug name and brand names in the prescription data. Dosage information and drug quantity was extracted using a regular expression that searched within the drug description. Combination therapies were excluded from our analysis.

We calculated maintenance dose by determining the average milligrams of drug per day for the last five prescriptions of each drug. This was done by calculating the total milligrams of drug administered for a single prescription divided by the number of days until the next prescription. We then averaged the milligrams of drug per day over the five most recent prescriptions. We excluded prescriptions with a dose quantity outside two standard deviations from the mean quantity across all participants for that drug. Subjects were required to receive a minimum of five prescriptions to be included in the analysis. We required drugs to have a minimum of 50 subjects with a maintenance dose to be included.

We divided the analysis of maintenance dose associations into three groups of drug-gene pairs. First, we investigated the relationship between drug-gene pairs that have an existing CPIC guideline. This indicates a strong level of evidence of a relationship between a drug and a gene. Second, we investigated drug-gene pairs which have some level of evidence in PharmGKB, but no existing CPIC guideline. These pairs still have some prior evidence indicating an association, but not enough to develop a dosage guideline. Third, we investigated all other drug-gene pairs where an interaction is indicated in DrugBank^12^. Data was grouped within each gene by predicted phenotype. For example, for *CYP2C9* participants were put into bins by metabolizer class (normal metabolizers (NM), intermediate metabolizers (IM), and poor metabolizers (PM). Phenotype groups with less than ten participants for a drug are excluded from analysis.

Association between maintenance dose and pharmacogenetic phenotypes was tested for 200 drugs using two types of non-parametric statistical association tests. We used both a Kruskal-Wallis one-way analysis of variance and Jonckheere-Terpstra trend tests to test for associations between each drug and gene pair. Both types of tests are necessary to detect various types of relationships between dosage and genetics. First, the Kruskal-Wallis test is used to identify any pharmacogenetic phenotype (e.g. CYP2C9 PMs) that have a significant difference in the dosage from other metabolizer classes. Second, Jonckheere-Terpstra tests for an ordered relationship in ranked groups. This is a natural fit for pharmacogenetic phenotypes since there is an inherent order in function which may lead to a linear relationship with dosage (e.g. NM > IM > PM). Resulting p-values are adjusted using a Bonferroni correction accounting for the multiple tests performed.

We tested the impact of the intronic *CYP2C19* variant rs3814637 on warfarin dose. We used a two-sided Jonckheere-Terpstra test on the allele dosage against the warfarin maintenance dose. Allele dosage was determined as the sum of the alternate alleles for rs3814637.

### Drug Side Effect Association with Pharmacogenetics

We tested the relationship between pharmacogenes and possible side effects for all drugs. For each drug included in the dosage analysis we identified all diagnoses in the primary care data in the year following the first exposure to the drug. Diagnosis codes in the primary care data are provided as Read Codes (version 2 and version 3), we mapped the Read Codes to ICD-10 codes including only the first three digits (the chapter and first two numerals). ICD-10 codes from chapters V, W, X, Y, and Z were excluded from analysis. Codes were required to have at least 100 events per drug to be included in the analysis.

Logistic regression was used to test the association between gene phenotypes and ICD-10 code incidence for each drug. This was set up using a binary indicator as the response variable and a one-hot encoding of gene phenotype. We included age (at time of first prescription), sex, genotyping array, and the first four principal components from a genotype PCA as covariates.

We evaluated three tiers of drug-gene relationship, as in the maintenance dose analysis. Drug-gene pairs with CPIC guidelines, drug-gene pairs with any level of evidence in PharmGKB but no CPIC guideline, and an exploratory analysis. For the exploratory analysis of side effect relationships we limited our search to drugs known to interact with *CYP2C9, CYP2C19*, and *CYP2D6*, as indicated by DrugBank. These genes were selected because they are promiscuous metabolizing enzymes with well defined pharmacogenetics.

## Results

The pharmacogenetic analyses presented here included a total of 201,498 participants after removing 20,615 participants not of european descent. More than 57 million prescriptions are contained within the primary care data, an average of 262 prescriptions per participant. Our initial drug list included 3,358 possible drugs. Of this, 200 were found in the UK Biobank prescription data with sufficient counts to be included in subsequent analysis.

### Dosage results

We sought to test the relationship between maintenance dose and pharmacogenes at a biobank scale. We performed this analysis using three groups of drug-gene pairs. Of the drugs with CPIC guidance for any of the nine genes queried, there were 22 that had the minimum of 50 participants for whom a maintenance dose could be calculated. We find that seven of the drug-gene pairs have a significant difference in the dosage across gene phenotypes (Kruskal-Wallis or Jonckeere-Terpstra *p* < 0.05, Table 1). We do not adjust for multiple tests because these are known relationships not discoveries. Warfarin and *CYP2C9* phenotypes had the most significant relationship (*p* ≅ 0, Jonckeere-Terpstra). The remaining twenty drug-gene pairs did not have a significant relationship between maintenance dose and gene phenotype.

**Table 1.**
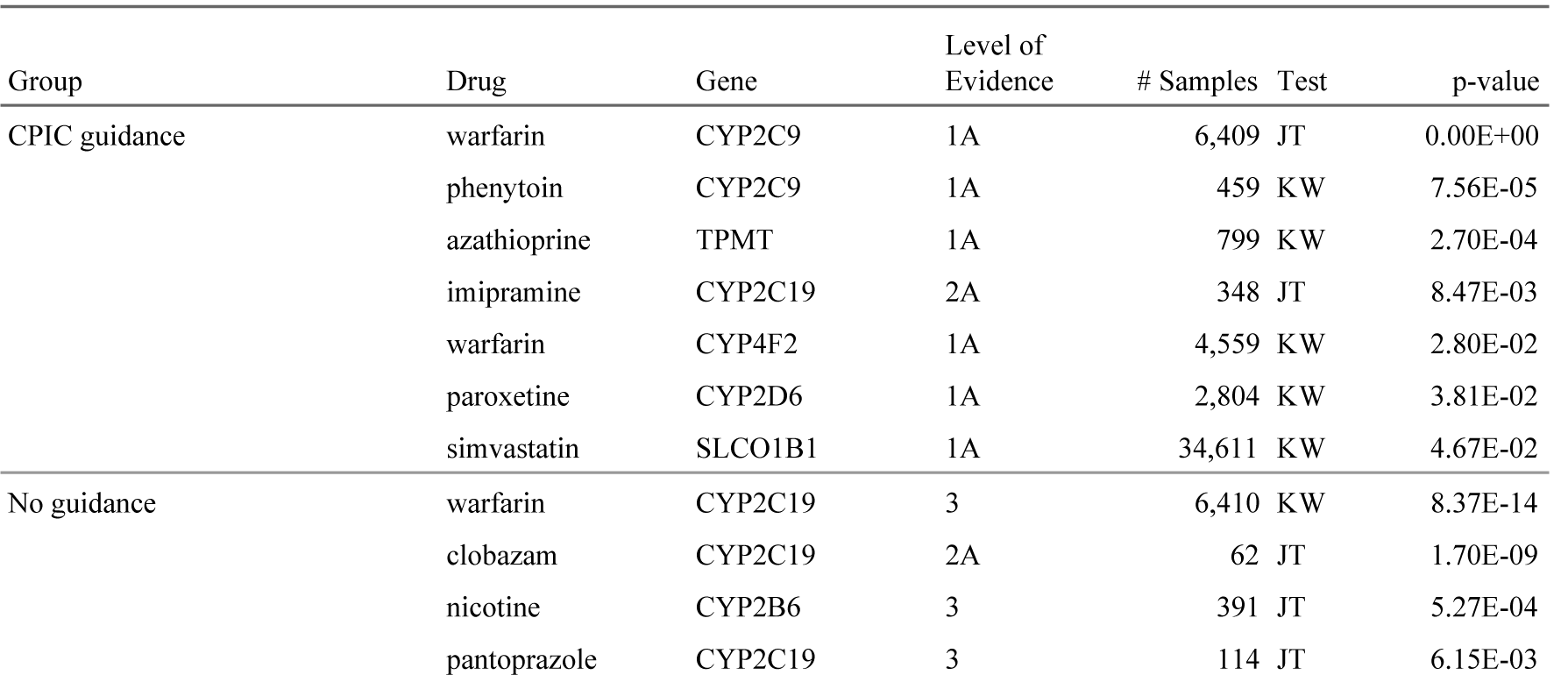
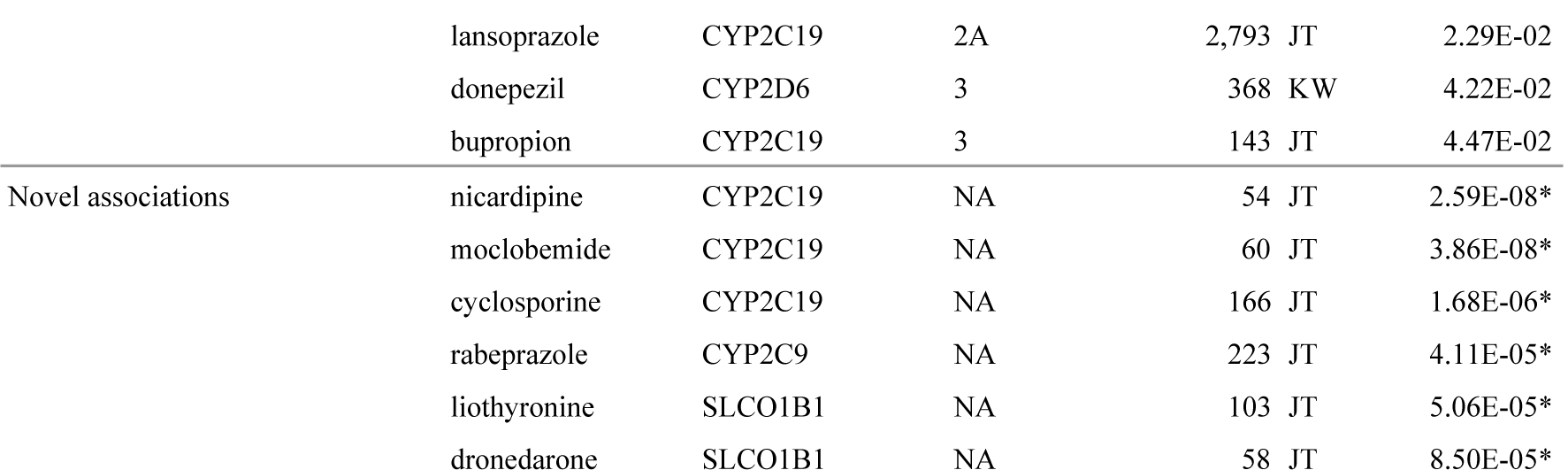
Drug-gene dose relationship results. Drug-gene pairs are presented in three groups: drugs with CPIC guidelines, without guidelines but PharmGKB evidence, and novel associations. Level of Evidence represents the maximum level of evidence for the drug-gene relationship in PharmGKB. p-values with a * are significant at *p* <= 8.6 × 10^−6^, bonferroni adjusted. Test indicates which type of test achieved the p-value shown (JT=Jonckheere-Terpstra, KW=Kruskal Wallis). Only results with a standard error less than 0.2 are included.

We then investigated association between maintenance dose and gene phenotype for drug-gene pairs with any level of evidence in PharmGKB but no CPIC guideline. We found seven drug-gene pairs with a p-value less than 0.05 for either the Kruskal-Wallis test or Joncheere-Terpstra trend test (Table 1). The most significant was the Kruskal-Wallis test for warfarin and CYP2C19 phenotype. Investigating the dose relationship with phenotype reveals that CYP2C19 normal metabolizers have a decreased maintenance dose compared to the other CYP2C19 metabolizer classes (Figure 1, second row, first column). We followed up on this finding by interrogating the association between rs3814637 and warfarin maintenance dose.

**Figure 1.**
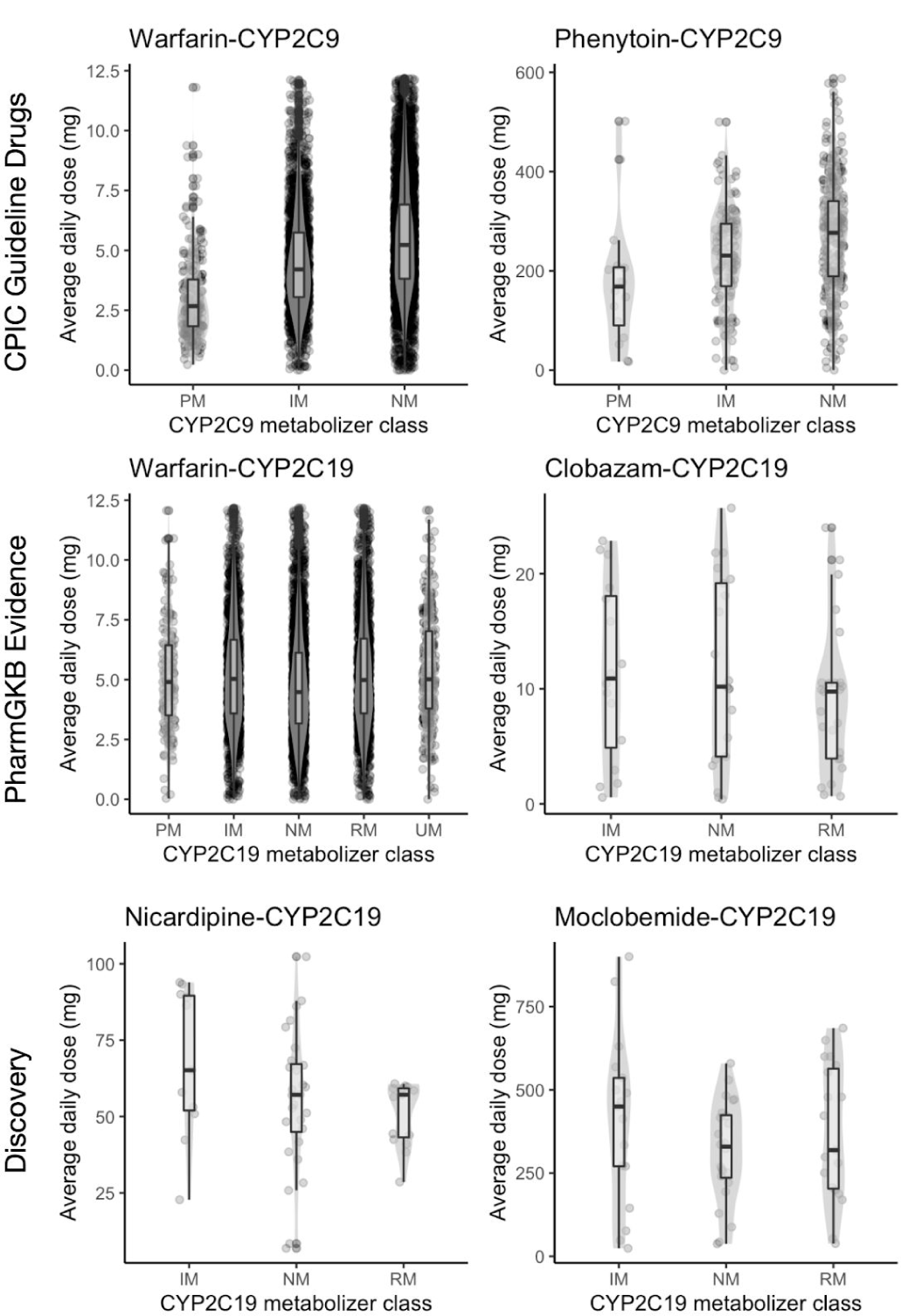
Box plots of maintenance dose for most significant drug-gene pairs. The top two most significant pairs are shown for each group. Enzyme metabolizer classes are represented along the x-axis and the distribution of maintenance dose along the y-axis.

The intronic variant rs3814637 within *CYP2C19* has been previously reported to be associated with warfarin response^13–15^. This variant is contained within several *CYP2C19* star alleles: *CYP2C19*1.004, CYP2C19*1.005*, and *CYP2C19*15.001*, all of which are normal functioning alleles. We observed that normal metabolizers had an average daily dose of 4.8 mg (compared to 5.3 mg for the other metabolizer classes). We then tested the association between rs3814637 and warfarin maintenance dose. We find a significant relationship between rs3814637 dosage and warfarin maintenance dose (*p* <= 1.0 × 10^−46^, two-sided Jonckheere-Terpstra, Fig. 2).

**Figure 2.**
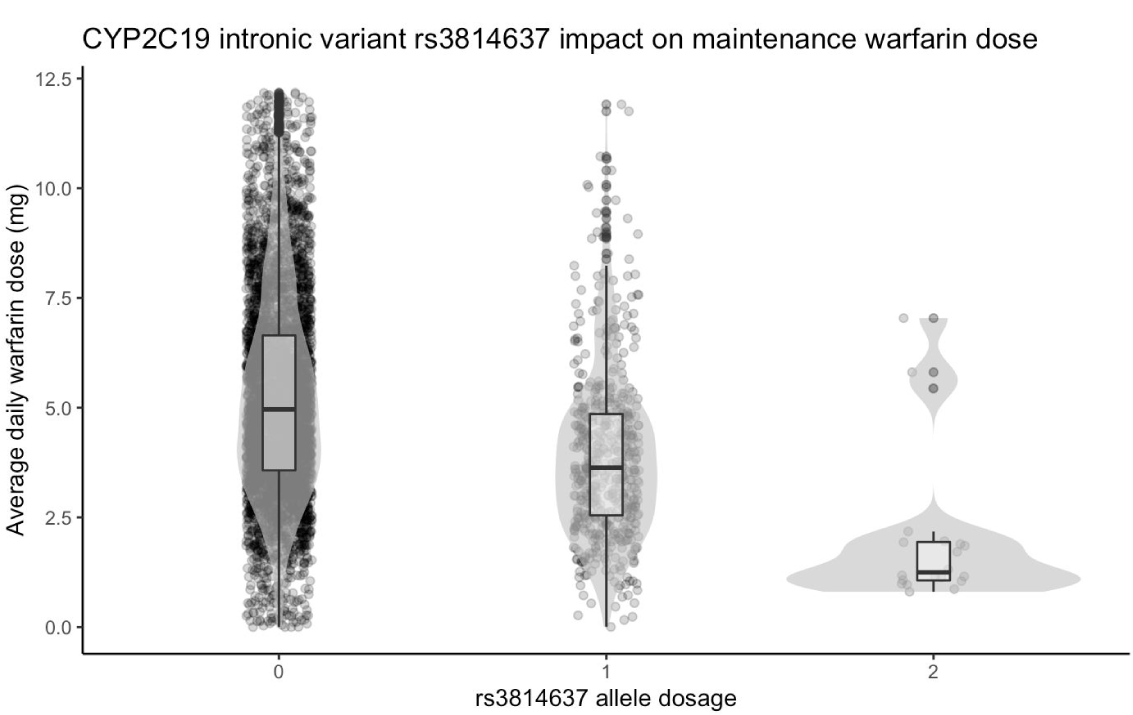
CYP2C19 intronic variant rs3814637 has a strong influence on warfarin maintenance dose. The x-axis indicates the alternate allele dosage (the sum of alternate alleles in each participant). The y-axis is the maintenance dose for each individual.

We then analyzed the relationship between maintenance dose and gene phenotype for drug-gene pairs that had no previous indication of a pharmacogenetic relationship but are known to interact. We tested 581 drug-gene pairs and found six significant relationships between dose and gene phenotype (*p* < 8.6 × 10^−6^, Jonckheere-Terpstra, bonferroni adjusted, table 3: Novel associations).

### Side effect results

We investigated the degree to which adverse drug reactions related to pharmacogenetics could be discovered by performing a statistical analysis of pharmacogene phenotypes and coded medical events within a one year window following the first administration of a drug. We again evaluated three drug-gene groups starting with drug-gene pairs with CPIC guidelines (Table 2, CPIC Guidance Group). The most significant side effect is a decreased incidence of herpes zoster diagnoses among CYP2C19 intermediate metabolizers on citalopram (*p* <= 8.76 × 10^−5^).

**Table 2.** Drug-gene side effect relationship results. Associations are presented in three groups: drug-gene pairs with CPIC guidelines, pairs with no guidelines but evidence in PharmGKB, and novel associations. Phenotype is the gene phenotype (IM: Intermediate Metabolizer, PM: Poor Metabolizer, RM: Rapid Metabolizer, UM: Ultrarapid Metabolizer, IF: Increased Function, PF: Poor Function). Odds ratio is the odds ratio relative to normal metabolizer or normal function alleles. * indicates significance with Bonferroni adjusted p-value threshold of 1.0 × 10^−5^. Only results with a standard error less than 0.2 are included.

| Group | Drug | Gene | Level of Evidence | Phenotype | ICD-10 | Code definition | Odds ratio | p-value |
| --- | --- | --- | --- | --- | --- | --- | --- | --- |
| CPIC Guidance | citalopram | CYP2C19 | 1A | IM | B02 | Herpes zoster | 0.53 | 8.76E-05 |
|  | simvastatin | SLCO1B1 | 1A | IF | M65 | Synovitis and tenosynovitis | 1.82 | 1.42E-04 |
|  | amitriptyline | CYP2C19 | 1A | RM | R53 | Malaise and fatigue | 1.55 | 1.74E-04 |
|  | amitriptyline | CYP2C19 | 1A | UM | J30 | Vasomotor and allergic rhinitis | 1.94 | 2.75E-04 |
|  | codeine | CYP2D6 | 1A | PM | A52 | Late syphilis | 1.78 | 3.30E-04 |
|  | ibuprofen | CYP2C9 | 1A | PM | E13 | Other specified diabetes mellitus | 2.00 | 4.90E-04 |
|  | clopidogrel | CYP2C19 | 1A | RM | B08 | Viral infections characterized by skin and mucous membrane lesions | 0.59 | 5.17E-04 |
|  | tamoxifen | CYP2D6 | 1A | IM | C50 | Malignant neoplasm of breast | 0.62 | 6.98E-04 |
|  | simvastatin | SLCO1B1 | 1A | PF | M79 | Unspecified soft tissue disorders | 1.49 | 7.46E-04 |
|  | simvastatin | SLCO1B1 | 1A | DF | M65 | Synovitis and tenosynovitis | 1.79 | 7.75E-04 |
| No Guidance | citalopram | CYP2D6 | 3 | IM | J45 | Asthma | 1.44 | 9.13E-05 |
|  | citalopram | CYP2D6 | 3 | IM | I50 | Heart failure | 1.56 | 1.12E-04 |
|  | simvastatin | CYP2C9 | 3 | PM | J01 | Acute sinusitis | 1.74 | 1.56E-04 |
|  | citalopram | CYP2D6 | 3 | IM | J64 | Unspecified pneumoconiosis | 1.56 | 5.74E-04 |
|  | propranolol | CYP2D6 | 4 | IM | O86 | Other puerperal infections | 1.85 | 6.38E-04 |
| Novel associations | diazepam | CYP2C9 | NA | PM | M19 | Osteoarthritis | 2.33 | 4.52E-06* |
|  | zopiclone | CYP2C9 | NA | IM | H91 | Unspecified hearing loss | 2.20 | 1.73E-05 |
|  | loratadine | CYP2D6 | NA | IM | M16 | Osteoarthritis of hip | 1.98 | 1.20E-04 |
|  | tramadol | CYP2B6 | NA | PM | H61 | Disorders of external ear | 1.95 | 1.86E-04 |
|  | quinine | SLCO1B1 | NA | IF | N39 | Disorders of urinary system | 1.95 | 1.87E-04 |

Next we looked to see if there are any side effects enriched among drug-gene pairs with any level of evidence but no CPIC guideline. The top five results are shown in Table 2 under “No Guidance”. We find several side effects of citalopram enriched among CYP2D6 intermediate metabolizers, including respiratory issues and heart failure. We also find an increased risk of sinus infections among CYP2C9 poor metabolizers on simvastatin, and an increased risk of puerperal infections among CYP2D6 intermediate metabolizers on propranolol.

We interrogated all other drugs known to be metabolized by CYP2C9, CYP2C19, or CYP2D6 for associations with side effects. This resulted in 4,806 independent association tests across 81 drugs. After multiple hypothesis corrections one side effect was significantly associated with a drug-gene pair: increased incidence of osteoarthritis in CYP2C9 poor metabolizers after taking diazepam. We show the top five results from the exploratory analysis in Table 2.

## Discussion

Biobanks offer a powerful solution for enabling the study of the relationship between drugs and genes. Large datasets linking genetic data and longitudinal clinical data are becoming more broadly available and allow interrogation of the relationship between drug response and pharmacogenetic phenotypes. Here we derived drug phenotypes in the form of maintenance dose and side effect incidence for more than 200,000 participants across 200 drugs in the UK Biobank and tested their association with well established pharmacogenetic phenotypes for nine genes.

While doctors may not be directly performing genetic tests to make treatment decisions, there is evidence that for some drugs genetics do influence the maintenance dose patients end up on through the natural clinical decision processes of drug selection or dose adjustment. We find evidence to support existing pharmacogenetic associations with maintenance dose. Among 22 drugs with CPIC guidance in our study we find evidence for a genetic influence on maintenance dose for seven drugs. For the remaining pairs with guidance, it is possible we are not likely to observe an association with maintenance dose because efficacy is difficult to measure or side effects leading to a dosage change are rare. Among drugs with any prior evidence of a pharmacogenetic relationship but no CPIC dosage guideline we find that maintenance dose supports the association for seven drug-gene pairs. Most notably, the *CYP2C19* intronic variant rs3814637 has a strong influence on warfarin maintenance dose. There is a gap in the amount of warfarin dosing variability that can be explained by genetics among individuals of African descent^16^. rs3814637 has nearly twice the allele frequency in the African population as it does in the European population (11.6% vs 6.7%)^17^. This may explain some of the missing heritability of warfarin response among Africans.

We discovered potential novel pharmacogenetic associations with maintenance dose for six drugs. All of these drug-gene relationships are known interactions (either the drug is metabolized or transported by the resulting protein). There is no literature supporting a pharmacogenetic relationship for five of the six pairs. However, moclobemide has an existing dosage guideline for *CYP2C19* genotype produced by Swissmedic, the Swiss agency for monitoring therapeutic products.

Our analysis of side effects reveals associations with side effects among drug-gene pairs. This analysis is limited due to the large number of tests requiring a strict multiple hypothesis testing threshold, but produces interesting hypotheses. At first glance many of the side effect associations seem unlikely, but literature evidence exists for many of the findings. For example, the most significant association among drugs with CPIC guidelines was a decreased incidence of herpes zoster among CYP2C19 intermediate metabolizers compared to CYP2C19 normal metabolizers treated with citalopram. However, two previous studies have demonstrated that SSRIs can lead to increased resistance to herpes zoster^18,19^. CYP2C19 intermediate metabolizers have an increased blood concentration of citalopram and may have an increased resistance to a herpes zoster infection. We also find CYP2C19 rapid metabolizers on clopidogrel have a decreased risk of viral skin lesions compared to CYP2C19 normal metabolizers. There is evidence that clopidogrel may inhibit viral clearance^20^. It may be possible that CYP2C19 rapid metabolizers have a lower concentration of clopidogrel and therefore the degree to which they are able to fight off viral infections is higher than that of CYP2C19 normal metabolizers. The most significant side effect association is between CYP2C9 poor metabolizers on diazepam having an increased incidence of osteoarthritis. There is no literature that suggests osteoarthritis may be a side effect of diazepam, although there are studies that suggest diazepam could be used to treat pain as a result of rheumatoid arthritis. Without further evidence it is difficult to say whether this relationship is a true side effect resulting from pharmacogenetics and not a correlation with the drug indication or some other statistical artifact.

This work has several limitations. First, we use pharmacogenetic alleles called from data imputed from genotyping arrays. We previously reported limitations in accuracy of the ability to accurately call alleles in several pharmacogenes from imputed data, notably in *CYP2D6*. The lack of structural variants in the dataset in addition to the inability to call rare variants may lead to inaccurate prediction of CYP2D6 phenotypes. Second, we broadly apply our maintenance dose algorithm to drugs in the UK Biobank. While this is effective for some drugs, better clinical end points may provide an improved representation of patient response. For example, a dose response curve may provide more fine grained insight into individual response and yield better insight into the genetics of drug response. It is challenging to broadly define response across drugs from numerous classes with varying indications and therapeutic indices. Even a single drug can be used for different indications and may require different doses to treat each indication. No catch-all definition will suffice, but maintenance dose does reveal insight into patient response. We also cannot for certain that the calculated maintenance dose is the true maintenance dose for a given patient. Third, the data we used to define drug usage is in the form of prescription orders. We do not know whether the prescriptions were filled or if the patient took the drug as prescribed. Finally, we do not provide any clinical validation of the predictions presented here; further followup is needed.

Biobanks are an immense resource that allow for pharmacogenetic association testing at an unprecedented scale. Longitudinal clinical data is critical to be able to define drug response phenotypes in order to accurately assess patient response to treatment and ultimately test genetic associations. As access to biobanks continue to expand and more data is available, the ability to perform pharmacogenetic studies at large scale will increase. We believe that these resources offer a promising avenue for discovery and will further advance the field of pharmacogenetics.

## Acknowledgments

G.M. is supported by the Big Data to Knowledge (BD2K) from the National Institutes of Health (T32 LM012409). R.B.A is supported by NIH/National Institute of General Medical Sciences PharmGKB resource (U24HG010615), NIH GM102365, and the Chan Zuckerberg Biohub. This research has been conducted using the UK Biobank Resource under Application Number 33722. We thank all the participants in the UK Biobank study. Most of the computing for this project was performed on the Sherlock cluster. We would like to thank Stanford University, the PharmGKB resource (NIH HG010615), and the Stanford Research Computing Center for providing the computational resources that contributed to these research results. Thank you to Adam Lavertu who helped develop the ideas that led to this work.

